# Characterization of an α-glucan-binding module from *Flavobacterium johnsoniae* as a founding member of carbohydrate-binding module family XXX

**DOI:** 10.64898/2026.01.30.702845

**Authors:** Tove Widén, Lauren S. McKee, Nicole Koropatkin, Johan Larsbrink

## Abstract

Carbohydrate-binding modules (CBMs) play crucial roles in carbohydrate-active enzymes by promoting substrate recognition and proximity, particularly for insoluble polysaccharides. Here, we report the discovery and characterization of a novel β-trefoil structured CBM associated with a GH87 α-1,3-glucanase from *Flavobacterium johnsoniae*, which accordingly was designated *Fj*CBMXXX_GH87_. The full-length enzyme efficiently hydrolyzed α-1,3-glucan (mutan) and α-1,3/α-1,6-glucan (alternan), whereas the catalytic domain alone displayed reduced activity, indicating that *Fj*CBMXXX_GH87_ enhances substrate interaction. Pull-down assays confirmed that *Fj*CBMXXX_GH87_ binds α-1,3-linked glucans, and structural investigation together with site-directed mutagenesis identified two distinct binding sites essential for protein-ligand interactions. Phylogenetic analysis showed that CBMXXX homologs are present together with enzymes from families GH87, GH13, GH16, and GH99, and potentially may comprise up to three binding sites. Together, these findings establish *Fj*CBMXXX_GH87_ as the founding member of a new CBM family, which may have broad functional versatility in polysaccharide recognition. This discovery expands the repertoire of β-trefoil CBMs and provides new insights into carbohydrate recognition strategies relevant to α-glucan degradation.

## Introduction

For carbohydrate-active enzymes (CAZymes), substrate binding is often modulated through the presence of carbohydrate binding modules (CBMs) which can be found appended to catalytic domains in multimodular proteins. The binding of an enzyme to its substrate at a suitable strength is essential to the enzyme’s functionality. Too weak binding limits the adsorption and thus the enzyme concentration close to the substrate, while an overly strong binding can limit the desorption from the substrate, causing a reduced turnover (1). This is especially true for insoluble substrates which cannot simply diffuse to the active site. In such cases, CBMs can also provide maintained enzyme proximity to the substrate even after dissociation, reducing the downtime of the enzyme until the next catalytic event. CBMs are classified and grouped into families in a sequence-similarity based manner, as are the catalytic CAZymes in the Carbohydrate-active enzymes database (CAZy, www.cazy.org, (2)).

CBMs can have different modes of binding, but most commonly their function is facilitated at least in part by the aromatic amino acids tryptophan, tyrosine, and, less commonly, phenylalanine (3). These can be arranged differently and, based on their configuration, CBMs may be divided into three types. In type A CBMs, the aromatic amino acids are placed in a planar configuration and with appropriate spacing to facilitate surface binding to flat crystalline substrates such as cellulose or chitin. In type B, they are arranged in a formation creating a groove where an internal segment of a polysaccharide chain can fit, creating an *endo*-type binding. Type C has shallower pockets facilitating binding to monosaccharides and/or terminal sugar moieties, enabling *exo*-type binding. Since the topography of the CBM needs to match the conformation of the target ligand, subtle differences in organization of the binding residues can vastly alter the CBM’s binding specificity, even though the overall structure of the protein may be very similar.

Some CBM domains of type B or C can have several binding sites, which may provide varying ligand specificity and enhanced affinity towards the target substrate (4). A type of CBM structure that has been shown to present several binding sites in one protein domain is the β-trefoil, characterized by the three-fold repeating symmetry of four β-strands, connected by loops and turns (5). Proteins with β-trefoil fold have thus far been found in three different CBM families: CBM13, CBM42 and CBM92, as well as in CBM-like lectins (2, 5). While these three families share a similar overall fold, their binding sites may be located or shaped differently, and a binding site may even be formed by amino acids from different regions of the polypeptide chain (6, 7). CBM13 modules have been found connected to many different kinds of CAZymes and shown to bind galactose, lactose, and xylan (6, 8) while proteins from CBM42 are most often associated with GH43 or GH54 enzymes and have been shown to bind arabinofuranosyl moieties on arabinoxylan and arabinogalactan (9, 10). CBM92 is a more recently discovered family associated with many diverse CAZyme families, such as GH5, GH30, and GH99 (7), and binding to β-1,3-glucans, β-1,6-glucans, and carrageenan has thus far been demonstrated in the family (11, 7, 12).

For well-studied polysaccharides such as cellulose or chitin, binding has been demonstrated by proteins in diverse CBM families (13). For less studied polysaccharides, there is however typically much less information regarding CBM-mediated binding. One such case is α-1,3-glucan, which is an insoluble polysaccharide found in the biofilm dental plaque and in the cell walls of some fungi (14, 15). In dental plaque, α-1,3-glucan is one of the main structural components and a virulence factor. It is produced from sucrose by transglycosylase enzymes secreted from oral bacteria such as *Streptococcus mutans* from where the polysaccharide gets its other name, mutan. α-1,3-glucan is important for the adherence of dental plaque to the teeth and enables the formation of the sticky extracellular matrix. Accumulation of plaque can eventually lead to the formation of anaerobic pockets where acidifying bacteria can thrive and cause erosion of the enamel of the teeth, leading to caries (14).

The only CBM family labeled with α-1,3-glucan-binding ability on the CAZy database is CBM24 (2), which encodes proteins typically found associated with α-1,3-glucanases (also known as mutanases). The first characterized CBM24 proteins were found appended to GH71 α-1,3-glucanases in *Trichoderma harzianum* and *Penicillium purpurogenum* (16). However, the mechanism for binding was not elucidated and important amino acid residues have not been identified. Suyotha et al. (17) studied the binding ability of non-catalytic protein domains in the GH87 α-1,3-glucanase Agl-KA from the bacterium *Niallia circulans*. They showed that two discoidin domains and a CBM6 domain independently bound weakly to α-1,3-glucan but that the three domains could bind more strongly together and that this binding increased the catalytic efficiency of the enzyme.

Though the prospect of using CAZymes for the degradation of dental plaque polysaccharides has been explored (18), an inherent challenge is the highly fluctuating environment with continuous saliva production, as well as food and beverages rapidly removing substances in the oral cavity, giving the enzymes limited time to act on the plaque (19). Discovery of new α-1,3-glucan-specific binding domains may therefore inspire useful tools to improve enzyme efficiency via improved adhesion to the substrate. Similarly, α-1,3-glucan-binding CBMs could be used to target fungal cell walls, for therapeutic purposes or selective labeling.

Recently a GH87 α-1,3-glucanase from *Flavobacterium* sp. EK-14, named Agl-EK14, was characterized by Takahashi et al. (20). The enzyme was shown to cleave insoluble α-1,3-glucan, both enzyme-synthesized and within fungal cell walls, and it was also shown through sequence analysis that the full protein comprised putative modules in addition to the catalytic domain. In this study we investigated a similar enzyme from *F. johnsoniae, Fj*GH87, and identified one of its non-catalytic modules to be a new type of β-trefoil CBM, the founding member of CBMXXX. The CBM domain, *Fj*CBMXXX_GH87_, was shown to bind insoluble α-1,3-glucans and different α-linked oligosaccharides, the connected GH87 module to be an active α-1,3-glucanase, and we further investigated the phylogeny of CBMXXX and its potential for multiple binding sites through bioinformatic and structure-function analyses.

## Methods

### Chemicals

α-1,3-glucan (mutan) and alternan (α-1,3/α-1,6-glucan) were synthesized enzymatically as described previously (21). Nigerooligosaccharides (α-1,3-linked glucoooligosaccharides, degree of polymerization 3-6) were purchased from Megazyme and nigeran (α-1,3/α-1,4-glucan) from Biosynth. All other chemicals were purchased from either Merck or Thermo Fisher Scientific unless noted otherwise.

### Molecular biology

Constructs for the target genes (locus tags: Fjoh_1208, Fjoh_3203) were amplified through PCR from genomic DNA from *F. johnsoniae* UW101 (ATCC 17061) using the primers listed in table S1. The PCR products for the respective proteins (see **Fig. 1** for *Fj*GH87 constructs; *Fj*GH13A for full-length Fjoh_1208 but lacking signal peptide and type IX secretion system C-terminal domain, *Fj*GH13A_CAT_ for its catalytic domain and *Fj*CBMXXX_GH13_ for its isolated CBMXXX domain) were cloned using InFusion cloning (Takara) into modified pET28A vectors containing a TEV protease cleavage site instead of the native thrombin site and modified T7 and TIR regions, as described previously (22, 23), and transformed into *E. coli* BL21 (λDE3) for protein production. Protein variants were created by site-specific mutagenesis by the QuickChange method (24). All constructs and mutations were verified through DNA sequencing (Eurofins Genomics).

**Figure 1.**
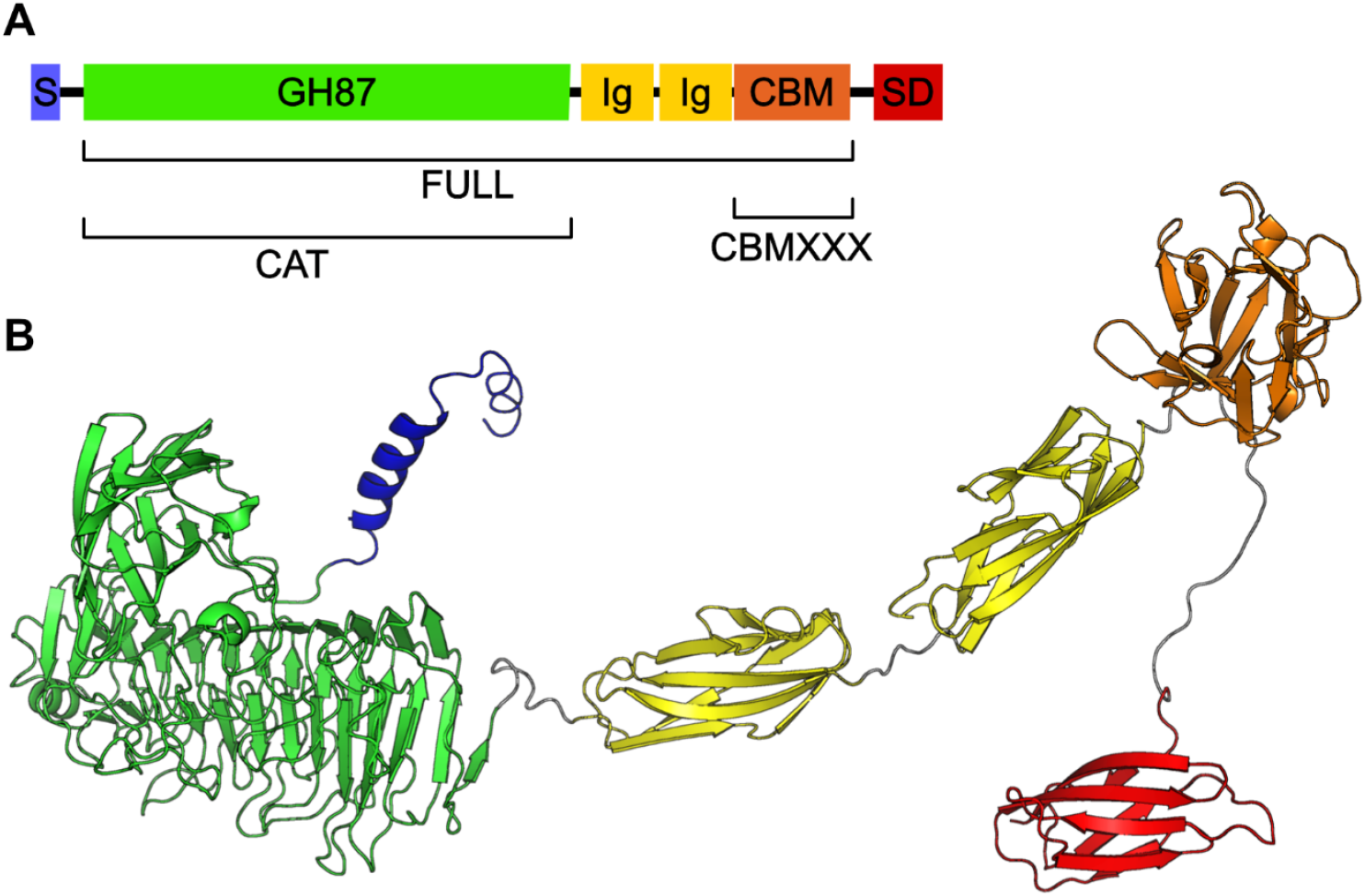
Domains of *Fj*GH87. **A**. Overview of constructs created. The whole protein length is 981 amino acids. FULL: The full-length protein but without signal peptide and C-terminal sorting domain (SD), 856 amino acids. CAT: only the catalytic GH87 domain, 550 amino acids. CBMXXX: The putative binding domain, 135 amino acids. Ig = immunoglobulin-like domains. **B**. AlphaFold3 model of *Fj*GH87, following the same color coding as in A, with linkers in gray.

### Protein production

For heterologous protein production, *E. coli* BL21 (λDE3) was cultured in lysogeny broth (LB) supplemented with antibiotics at 37 °C and 200 rpm shaking until an OD_600_ of ∼0.5 was reached, after which expression was induced through the addition of isopropyl β-D-1-thiogalactopyranoside (IPTG) to a final concentration of 0.25 mM, after which the cells were incubated at 16 °C overnight.

The cells were harvested by centrifugation (5000 ×g 10 min), resuspended in 20 mM tris(hydroxymethyl)aminomethane (Tris) buffer, pH 8, containing 250 mM NaCl, and disrupted by sonication. Cell debris was removed by centrifugation (18,000 ×g, 10 min), and proteins were purified using immobilized metal ion affinity chromatography on an ÄKTA system (GE healthcare) with 5 mL HisTrap™ Excel columns, using 50 mM Tris, pH 8, with 250 mM NaCl as binding buffer and a linear gradient of the same buffer containing 250 mM imidazole for elution. The protein samples were concentrated by ultrafiltration (Amicon Ultra-15, Merck-Millipore).

Sodium dodecyl sulfate polyacrylamide gel electrophoresis (SDS-PAGE) using Mini-PROTEAN TGX Stain-Free Gels (BIO-RAD) was used to verify protein purity and protein concentrations were measured using a Nanodrop 2000 Spectrophometer (Thermo Fisher Scientific) and extinction coefficients predicted using Benchling.

### Enzyme assays

Screening of enzyme activities was conducted in 150 µl reactions containing 50 mM sodium acetate buffer at pH 5, 1 µM enzyme, and 1 mg/ml of either of the polysaccharides α-1,3-glucan, alternan, dextran, starch, cellulose (Avicel), beechwood xylan, α-chitin, β-chitin or yeast α-mannan, or 100 µM of the oligosaccharides nigerose, nigerotriose, nigerotetraose, nigeropentaose, nigerohexaose, maltohexaose, isomaltohexaose, or cellohexaose, respectively. The reactions were incubated at 25 °C with 1000 rpm shaking in a thermomixer (Eppendorf) for 24 h and then stopped through heat inactivation at 85 °C for 2 min. The reactions were then centrifuged to remove residual polysaccharides and precipitated proteins and the supernatants further analyzed using high-performance anion-exchange chromatography with pulsed amperometric detection (HPAEC-PAD) on a Dionex IC-5000+ (Thermo Fischer Scientific) equipped with a Dionex™ CarboPac™ PA200 IC Column (Thermo Fischer Scientific), as described earlier (21). Standards with glucose, nigerose, nigerotriose, nigerotetraose and nigeropentaose were used for identification and quantification of reaction products.

### Binding studies

Binding of CBMs to the insoluble polysaccharides α-1,3-glucan, alternan, nigeran, starch (potato), cellulose (Avicel), wheat arabinoxylan, chitin (from shrimp shells) and β-glucan (from yeast) were determined through pull-down assays. Suspensions of 100 µL were prepared with final concentrations of 0.3 mg/mL *Fj*CBMXXX_GH87_ and 1 mg/mL polysaccharide in buffer (50 mM Tris, pH8, 250 mM NaCl). The suspensions were vortexed and incubated at 25 °C and 1000 rpm shaking for 1 h, after which they were centrifuged and the proportion of unbound protein determined through SDS-PAGE.

Binding to soluble saccharides was assessed through isothermal titration calorimetry (ITC). Initial testing, including an investigation of *Fj*CBM26_GH13_, was performed using a TA Instruments standard volume NanoITC. For each experiment, 1.3 mL of 60 μM protein was added to the sample cell and the reference cell was filled with distilled water. The sample injection syringe was loaded with 250 μL of the appropriate ligand concentration (0.5–3 mM), aiming to fully saturate the protein by the end of 25 injections of 10 μL. Titrations were performed at 25 °C with a stirring speed of 250 rpm.

All other experiments were performed using a MicroCal iTC_200_. For each experiment 300 μl of 25 μM protein was added to the sample cell and the reference cell was filled with buffer (50 mM Tris, pH8, 250 mM NaCl). The sample injection syringe was loaded with 40 μL of the appropriate ligand concentration (0.5–4 mM), aiming to fully saturate the protein by the end of 20 injections of 2 μL. Titrations were performed at 25 °C with a stirring speed of 750 rpm.

### Circular Dichroism

To verify that the folding of the protein variants created was similar to that of the wild type *Fj*CBMXXX_GH87_ protein, the proteins were analyzed by circular dichroism (CD). The proteins were buffer exchanged into 10 mM sodium phosphate, pH 8. Measurements were performed in 300 µL samples, including 2 µM protein, in a 2 mm pathlength quartz cuvette. The CD spectrum of each protein was recorded from 190 to 240 nm in 1 nm steps using a J-815 CD spectrometer (Jasco).

### Bioinformatic analyses

The *Fj*CBMXXX_GH87_ protein sequence was used as a query in a BLAST search against the ClusteredNR database (25). The 250 top hits of sequences aligning to *Fj*CBMXXX_GH87_ were collected and re-aligned using Clustal Omega (26) and a phylogenetic tree constructed using iqtree (27), using default settings including choice of optimal substitution model (WAG+G4) and 1000 ultrafast bootstraps (28). The complete protein sequences of the hits were then submitted to dbCAN3 HMMER to identify other potential CAZy domains connected to the identified CBMXXX modules. The CBMXXX phylogenetic tree, including the domain annotations for multimodular proteins, was visualized with iTOL (29).

## Results and Discussion

### GH87 characterization

A single GH87 module has been identified in the *F. johnsoniae* UW101 genome, and it was named *Fj*GH87. The full protein comprises 981 amino acid residues, and domain analysis shows a striking resemblance to Agl-EK14 from *Flavobacterium* sp. EK-14 (20) (84 % sequence identity), with an N-terminal signal peptide, the GH87 domain, followed by two immunoglobulin-like (Ig-like) domains, a Ricin B-like domain, and a C-terminal type IX secretion system sorting tag (**Fig. 1**) (30). An AlphaFold3 prediction of the protein structure validated the sequence-based domain annotation (31), and the predicted structure was used to guide cloning for heterologous expression in *E. coli*. The enzyme was cloned as a full-length construct (signal peptide and C-terminal sorting tag removed) and as the single GH87 domain. The activity of the full length *Fj*GH87 enzyme was screened on various substrates, and activity could be seen on the polysaccharides α-1,3-glucan, alternan (mixed-linkage α-1,3/1,6-glucan; **Fig. S1**) and weakly on starch but not on dextran (α-1,6-glucan), cellulose (Avicel), beechwood xylan, α-chitin, β-chitin, or yeast α-mannan. On oligosaccharides, the enzyme showed some activity on maltohexaose but not on isomaltohexaose or cellohexaose, demonstrating a clear preference for α-1,3-glucosidic linkages. When assayed on other α-1,3-linked oligosaccharides (nigerooligosaccharides), in addition to nigerohexaose, the enzyme also cleaved nigeropentaose and nigerotetraose, but not nigerotriose or nigerose, indicating that the minimal substrate is a tetrasaccharide. From nigerohexaose, the main end-point product was nigerotriose, but also small amounts of nigerose were formed (**Fig. S2**). From nigeropentaose, the end-point products were nigerotriose and nigerose, and from nigerotetraose, a mixture of nigerotriose, nigerose and glucose was produced.

The recent studies of Agl-EK14 were conducted using a full-length construct, including the Ig-like and Ricin B-like domains, but whether these additional domains contribute to its activity was not investigated (20). When we compared the activity of the full-length *Fj*GH87 enzyme to that of the lone catalytic domain, on α-1,3-glucan using equimolar amount of enzyme over four hours (**Fig. 2**), the full-length enzyme produced almost double amounts of the final products glucose, nigerose and nigerotriose compared to the sole catalytic domain. For both enzymes, nigerotetraose was produced as an initial product that was then gradually further degraded, but the amounts of both initial nigerotetraose and the later removal was greater for the full-length enzyme, i.e. higher initial amount and lower final amount. Ig-like fold domains have been found in various proteins (32, 33), and while these could potentially be involved in carbohydrate-binding, we hypothesized that the difference in activity was more likely due to the uncharacterized Ricin B-like domain. A structural prediction revealed this domain to form a β-trefoil domain similar to CBM13, CBM42 and CBM92 proteins, which could represent a novel CBM family facilitating binding to α-1,3-glucan.

**Figure 2.**
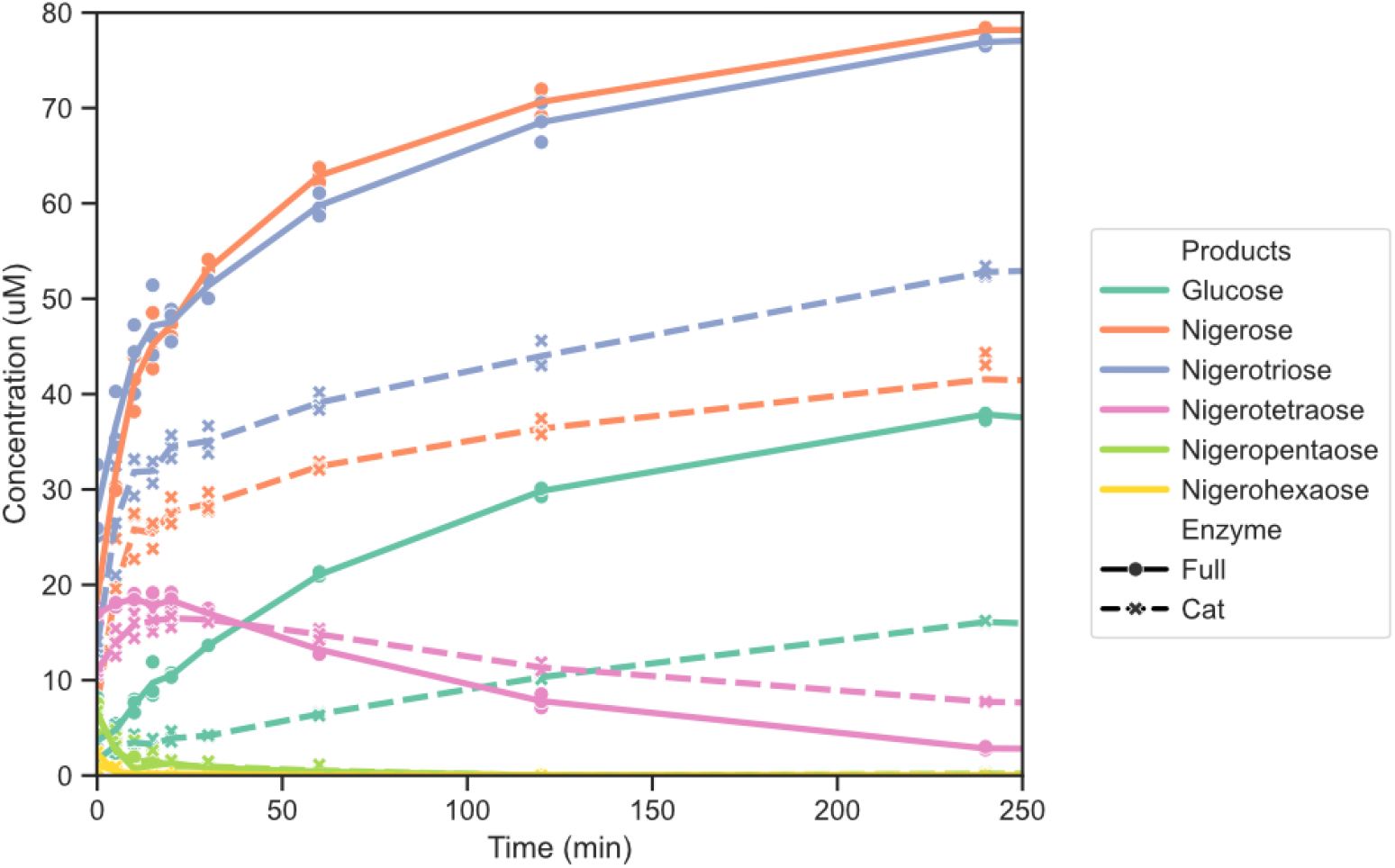
Hydrolysis reactions by *Fj*GH87. Comparison of the enzymatic activities of the full-length *Fj*GH87 (solid lines) and the lone catalytic domain of *Fj*GH87 (dashed lines). The release of glucose and nigerooligosaccharides are shown as individual data points, with lines connecting the average values.

### Binding to insoluble polysaccharides and preliminary structural analysis of FjCBMXXX_GH87_

To investigate the potential binding of the β-trefoil domain, from now on referred to as *Fj*CBMXXX_GH87_, it was produced as a separate construct and its binding abilities was assessed in pull-down assays (**Fig. 3**). The protein was confirmed to bind α-1,3-glucan and alternan (α-1,3/α-1,6-glucan), potentially nigeran (α-1,3/α-1,4-glucan) to a small extent, but not starch, cellulose, wheat arabinoxylan, chitin, or yeast β-glucan.

**Figure 3.**
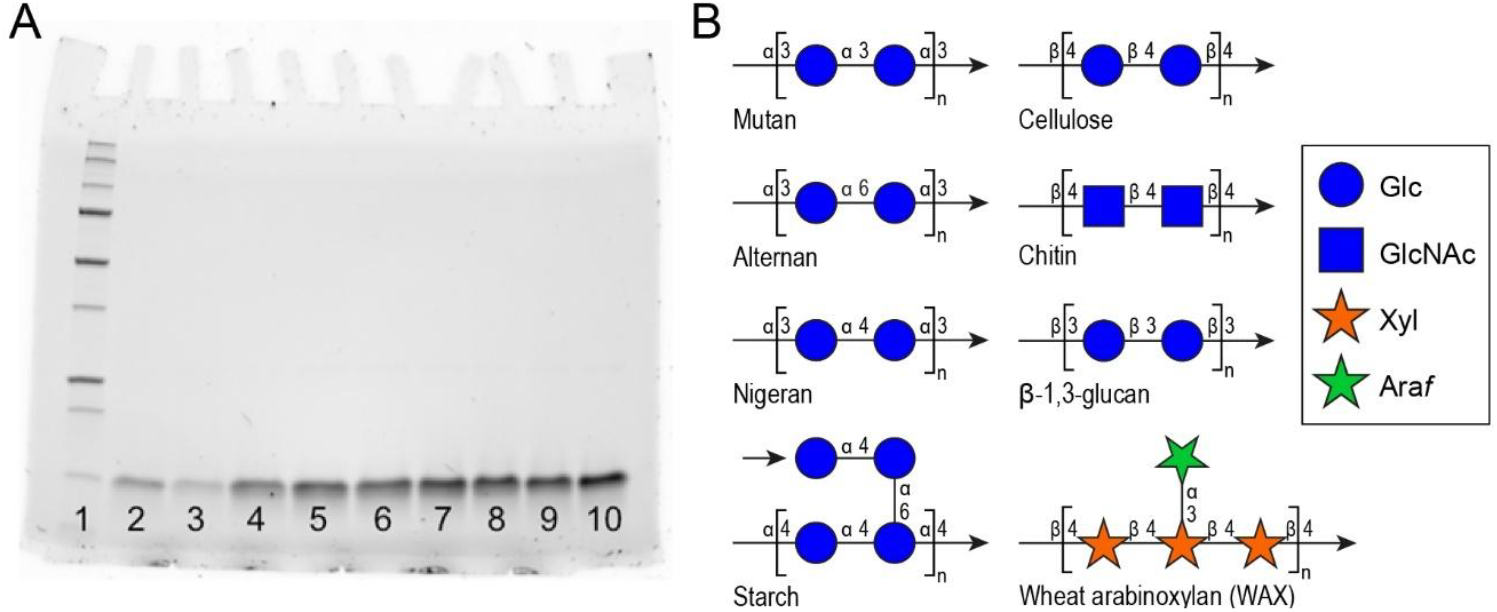
**A**. Pull-down assay of *Fj*CBMXXX_GH87_, where the protein was incubated with different insoluble polysaccharides, followed by SDS-PAGE analysis of protein remaining in solution. 1: molecular weight ladder, 2: α-1,3-glucan, 3: alternan, 4: nigeran, 5: starch, 6: cellulose, 7: wheat arabinoxylan, 8: chitin, 9: yeast β-glucan, 10: negative control. The lower concentration of protein compared to the control indicates *Fj*CBMXXX_GH87_ is bound to the polysaccharides α-1,3-glucan and alternan, but not significantly to nigeran, starch, cellulose, wheat arabinoxylan, chitin or yeast β-glucan. **B**. Representative composition and linkages of the tested polysaccharides, following the symbol nomenclature for glycans (SNFG, (34)).

Proteins with a β-trefoil fold have a threefold symmetry, where the different parts, or leaflets, can be referred to as the α, β, and γ sub-domains. In other β-trefoil fold CBM families, one or more of these leaflets may contain carbohydrate-binding sites, with a maximum of three independent binding sites possible (7, 10). Analysis of the AlphaFold model of *Fj*CBMXXX_GH87_ showed the three expected leaflets, which were named in order of occurrence counting from the N-to the C-terminus (**Fig. 4**). In both the β- and γ leaflets, we could observe exposed tryptophan residues that we speculated could be involved in the glycan-binding ability of the protein, and these positions on the proteins were referred to as the β-and γ-sites, respectively. The corresponding α-site did not contain any aromatic amino acids in equivalent positions. The α-site contains two surface-exposed tyrosine residues, but these however appear stacked without enough space between them to facilitate binding of a carbohydrate moiety, and we hypothesized that this is not a binding site. Possibly, the lack of an α-site could be explained by the position of this site which would be in close proximity to the domain’s N- and C-termini, i.e. the attachment site to the rest of *Fj*GH87. A binding site located on the α leaflet could then likely be blocked by the full-length enzyme’s other domains.

**Figure 4.**
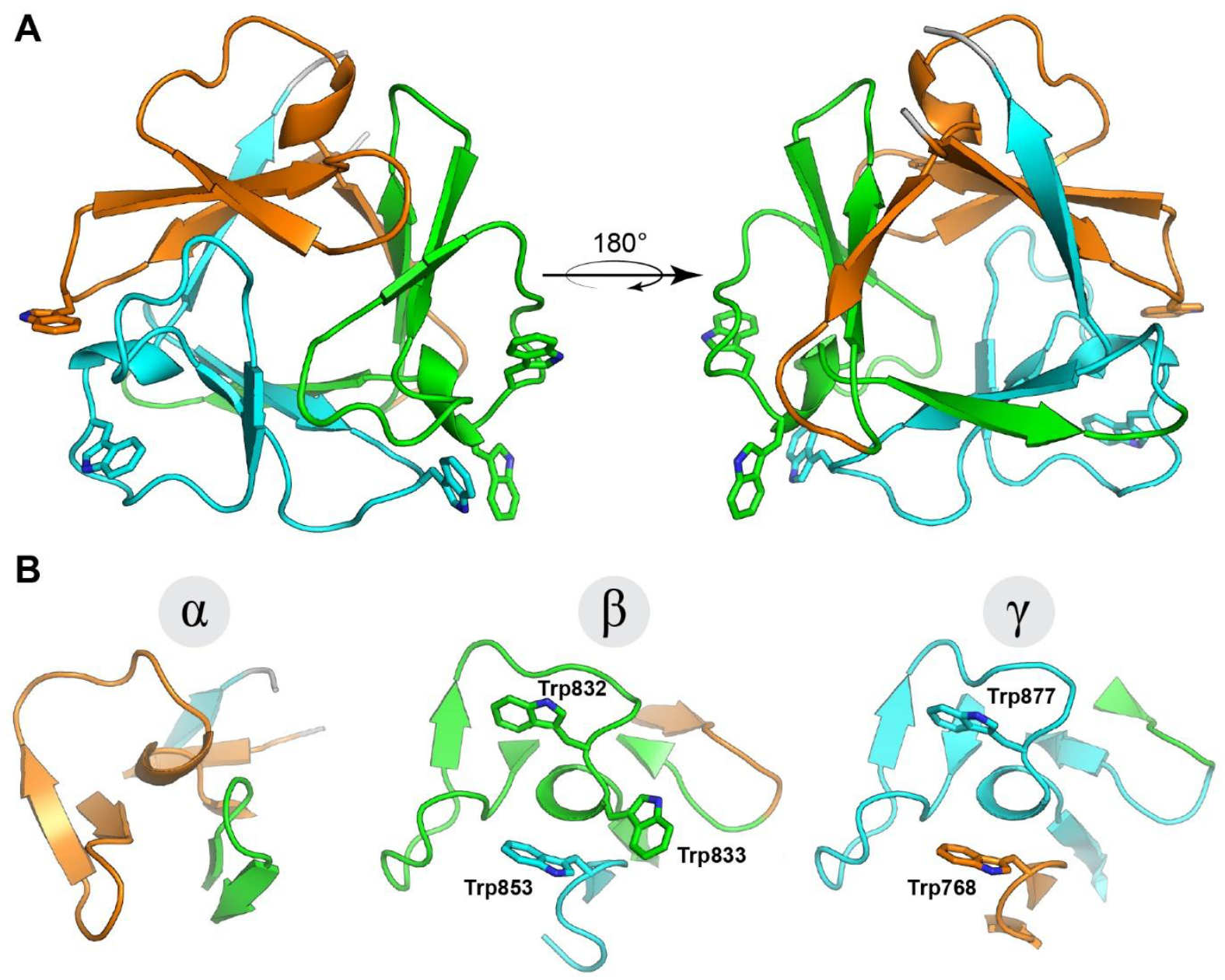
Predicted structure of *Fj*CBMXXX_GH87_ and comparison of leaflets. **A**. The AlphaFold-predicted structure of *Fj*CBMXXX_GH87_ with putative binding residues shown. The leaflets are color-coded based on their sub-domain designation: α-leaflet – orange (originating from the N-terminal end), β-leaflet – green, and γ-leaflet – cyan (ending at the C-terminal end). **B**. The leaflets of *Fj*CBMXXX_GH87_ aligned to each other, with putative binding residues shown and labelled. The α-leaflet lacks any apparent binding residues while the β- and γ-leaflets contain three and two putatively binding tryptophan residues, respectively.

The putative β- and γ-sites include three and two tryptophan residues, respectively; β: W832, W833 and W853, and γ: W768 and W877 (numbered by their position in the full-length protein; **Fig. 4B**). To assess the importance of these residues we converted them to alanine using site-directed mutagenesis of the encoding plasmid. We also substituted the two tyrosine residues close to the position of the would-be α-site to alanine, though these were as predicted not involved in binding (S1-2). To investigate whether the substitutions had any significant impact on the folding of the variants, they and the wild type *Fj*CBMXXX_GH87_ protein were subjected to circular dichroism, which showed highly similar signals for all proteins, indicating no detrimental effects from the substitutions (**Fig. S3)**. The W832A and W833A substitutions or either of W853A, W768A or W877A, as well as complete substitution of all tryptophans in either binding site, resulted in clear reduction of the binding ability on both α-1,3-glucan and alternan, as assesed by pull down assays (**Fig. S4-6**), indicating that both binding sites are necessary for sufficiently strong binding to be detected via this method. Having verified the identity of the binding residues, we also wanted to assess their importance and specificity quantitatively, and also investigate whether the degree of polymerisation of the saccharide affected the affinity. We therefore moved on to study the binding to soluble oligosaccharides.

### Binding to oligosaccharides

The binding of *Fj*CBMXXX_GH87_ to soluble saccharides was assessed using quantitative isothermal titration calorimetry (ITC); the dissociation constants (K_D_) are displayed in **Table 1** and representative traces in **Fig. S7**. Binding of *Fj*CBMXXX_GH87_ was detected to malto-, isomalto- and nigerooligosaccharides but not to glucose, β-cyclodextrin, or dextran. The affinity was not consistently higher with increasing degree of polymerization but varied with the glycosidic linkage. While no binding could be detected to glucose, the K_D_ values for nigerooligosaccharides decreased with increasing length, from 1420 µM for nigerose, to 826 µM for nigerotriose and 437 µM for nigeropentaose. In contrast, the K_D_ values for isomaltotriose were higher than for isomaltose, at 971 compared to 662 µM. Lastly, the K_D_ value for maltose was 609 µM, and decreased for maltotriose and maltopentaose to 259 and 292 µM, respectively. It is somewhat surprising, in the context of the above-mentioned pull-down assays, that *Fj*CBMXXX_GH87_ bound more strongly to maltose and isomaltose than nigerose, since it was the α-1,3-linked polysaccharides that showed the most obvious binding in those previous assays. It is also interesting that the affinity does not follow a clear pattern when it comes to degree of polymerization. This could indicate that while the protein could bind to several smaller oligosaccharides relatively independent of linkage, longer oligosaccharides need to be accommodated more specifically, resulting in a narrower specificity the longer the saccharide gets. This could also explain why no binding to either dextran or cyclodextrin was observed, despite them including the same kind of linkages that were recognized for di- and trisaccharides.

**Table 1.**
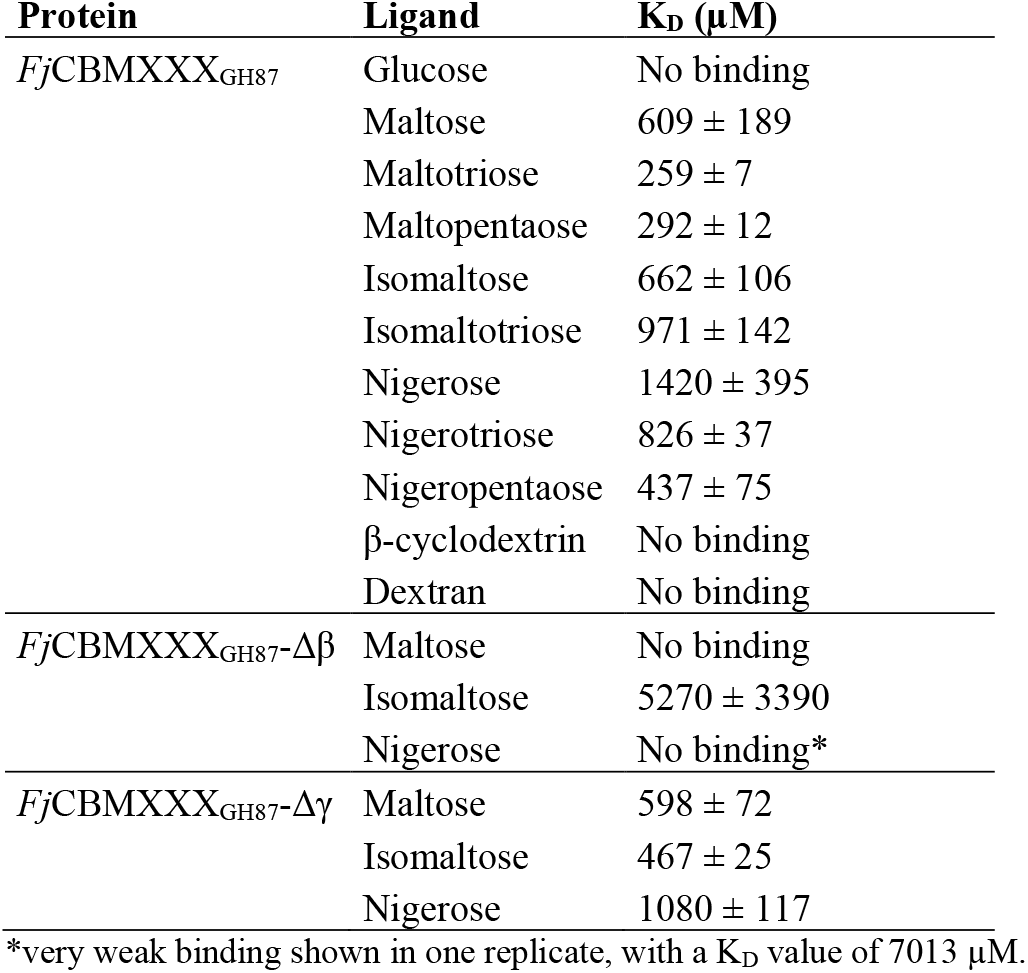
Dissociation constants of *Fj*CBMXXX_GH87_ and variants. Binding studies were performed in duplicate experiments using ITC.

To investigate the binding site variants, *Fj*CBMXXX_GH87_-Δβ (W832A, W833A, W853A) and *Fj*CBMXXX_GH87_-Δγ (W768A, W877A) were subjected to ITC with the same oligosaccharides, **Table 1** and **Fig. S8**. The binding of *Fj*CBMXXX_GH87_-Δβ variant was severely impacted for all the tested oligosaccharides, with either no reliable binding, or very high K_D_ values, over 5 mM for isomaltose and around 7 mM for one of the nigerose replicates. By contrast, the binding of *Fj*CBMXXX_GH87_-Δγ variant was not markedly impacted compared to the wild type, with similar or slightly lower K_D_ values for the disaccharides, 598 compared to 609 µM for maltose, 467 compared to 662 µM for isomaltose, and 1080 compared to 1420 µM for nigerose.

Taken together, the results indicate the dominance of the β-site over the γ-site in conferring binding. This is somewhat contradictory to the results from the pull-down assays which showed that alteration of any of the putative binding residues severely impacted the proteins’ ability to bind to both the α-1,3-glucan and alternan polysaccharides. A plausible explanation is that the γ-site does participate in binding, though with considerably lower affinity than the β-site, making its contribution undetectable by ITC. However, when binding to polysaccharides, both sites may be required to achieve sufficient binding strength for the protein to sediment into the pellet. Possibly, there are certain conformations that are more prevalent in the polymeric form of the ligands, not sampled in these experiments with mono- and oligosaccharides, that better suit the γ-site.

### Phylogeny and features of CBMXXX

To investigate the diversity and distribution of CBMXXX modules, and to elucidate which CAZymes these modules are commonly associated with, we performed a BLAST search against the ClusteredNR database with *Fj*CBMXXX_GH87_ as query. CBMXXX appears to be a relatively small family, exclusively found in bacteria, and the BLAST analysis yielded 376 sequences with a query coverage over 60 % and percent identity over 30 %. The 250 top hits were used to construct a phylogenetic tree (**Fig. 5**). By coloring the branches based on the phylum from which each protein originates, a wide diversity of CBMXXX-encoding organisms could be found, including many from Bacteroidota, Pseudomonadota, Bacillota, Actinomycetota and Myxococcota.

**Figure 5.**
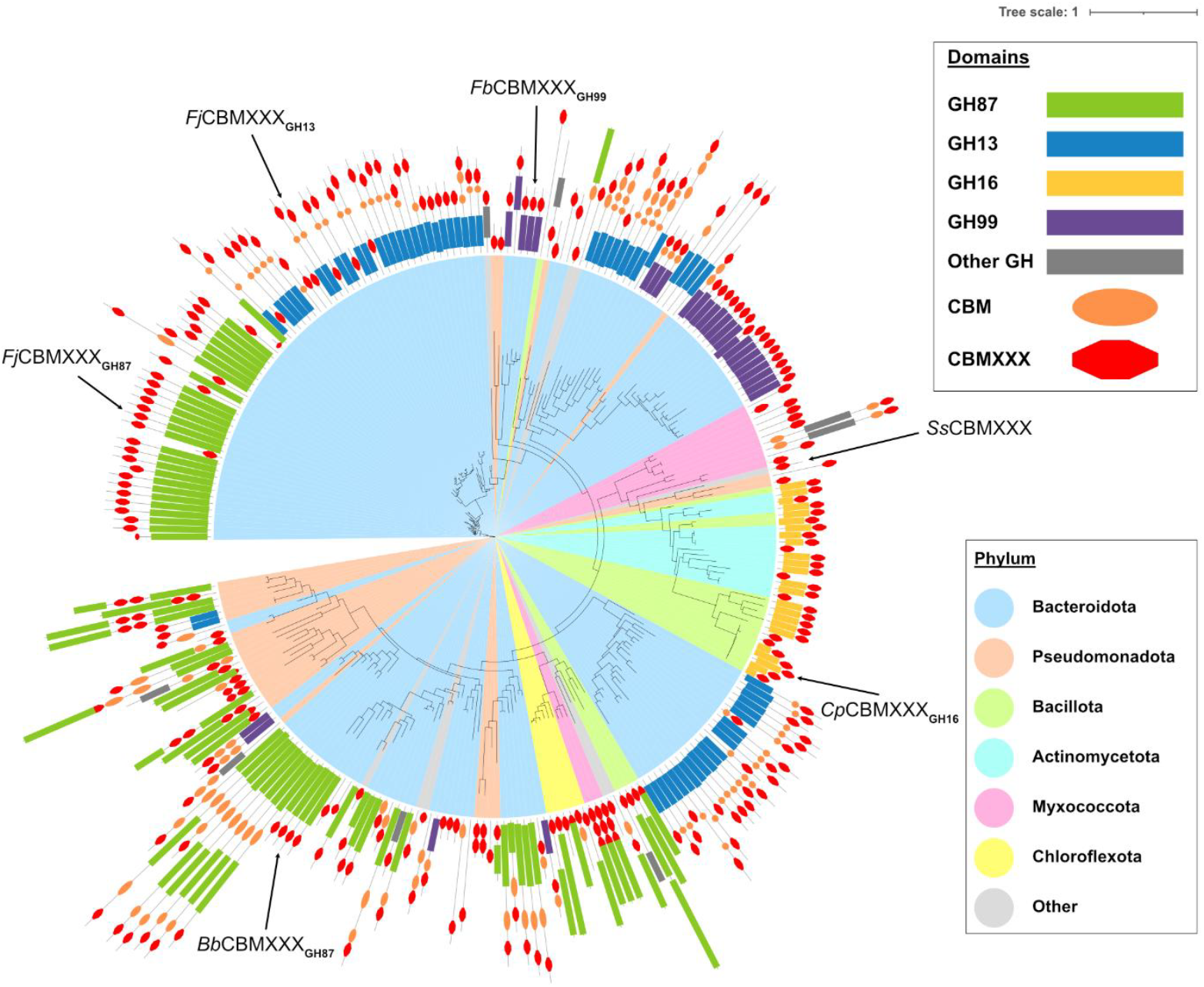
Phylogenetic tree of CBMXXX. *Fj*CBMXXX_GH87_ and other proteins shown in Fig. 6, *Ss*CBMXXX, *Cp*CBMXXX_GH16_, *Bb*CBMXXX_GH87_, *Fv*CBMXXX_GH99_ and *Fj*CBMXXX_GH13_ are indicated with arrows. The outer ring shows the modularity of the full-length proteins, with red diamonds indicating CBMXXX modules, differently colored rectangles indicating glycoside hydrolases (GH87 green, GH13 blue, GH99 purple, GH16 yellow, others gray), and orange ovals showing non-CBMXXX CBM families. A complete version including all GenBank IDs is shown in Fig. S9.

Graphical depictions of the modularity of the corresponding full-length proteins the CBMXXX modules were included as an added layer of information in the tree, which showed that CBMXXX is most often associated with GH87, as in *Fj*GH87. Other families that were commonly found fused to CBMXXX domains were GH13 (mainly amylases), GH99 (mainly α-1,2-mannosidases), and GH16 (various endo-acting β-glycanases). We could observe that especially the CBMXXX domains connected to GH16 or GH99 enzymes clustered together in the tree, indicating a similar evolutionary origin. CBMXXX domains appended to GH87 or GH13 enzymes also clustered together, but not in a single clade, and possibly, co-occurrence of CBMXXX with these catalytic partners may have arisen on multiple occasions. A few single CBMXXX domains were also observed that were not part of larger CAZymes, and these proteins may potentially either act as lectins or be a result of low-quality sequencing leading to fragmented genes.

Interestingly, the sequence analysis revealed one more CBMXXX-containing protein encoded by *F. johnsoniae*, Fjoh_1208 (Genbank ID WP_187775473.1), which is comprised of a GH13 domain followed by a CBM26 module and a CBMXXX-domain (*Fj*CBMXXX_GH13_). Since many CBMXXX domains were found together with GH13 enzymes, we chose to study this protein as well. The *Fj*CBMXXX_GH13_-domain has 60 % sequence identity to *Fj*CBMXXX_GH87_ and is thus not expected to be a result of a recent gene duplication. Structural comparisons using AlphaFold models revealed similar binding sites, but with a difference in the β-site in *Fj*CBMXXX_GH13_, which has a phenylalanine in the corresponding position of W332 in *Fj*CBMXXX_GH87_. The γ-site in *Fj*CMBXXX_GH13_ is similar to the β-site in *Fj*CBMXXX_GH87_, with two tryptophans and a tyrosine residue. We were unable to quantify binding of *Fj*CBMXXX_GH13_ in either pull-down assays or ITC measurements to a wide range of insoluble and soluble glycans. However, we could observe very weak (not quantifiable) binding using ITC to maltopentaose (**Fig. S10**). Possibly, the slight difference in the binding sites compared to the dominant β-site in *Fj*CBMXXX_GH87_ is a reason for this poor binding, or that we have yet to test the actual target carbohydrate for this particular protein. The connected GH13 enzyme was shown to be active on maltooligosaccharides and the other binding module in the full-length protein, from CBM26, was shown to bind to maltooligosaccharides (**Fig. S11**.), which demonstrates that the overall protein is functional.

Of the GH16 enzymes connected to CBMXXX domains, all belong to subfamily 24 (GH16_24), from which there is only one characterized member to date, the mucin endo-β-1,4-galactosidase Endo-β-Gal_GnGa_ from *Clostridium perfringens* (35). The crystal structure of this enzyme (PDB: 1UPS) also includes its CBMXXX domain but this protein domain has not been studied functionally. This CBMXXX-domain, hereafter called *Cp*CBMXXX_GH16_, has 35 % sequence identity to *Fj*CBMXXX_GH87_. As in *Fj*CBMXXX_87_, the α-site of *Cp*CBMXXX_GH16_ is seemingly lacking binding residues, while the β-site contains the identically positioned three tryptophan residues. The γ-site is, however, somewhat different. Where the site in *Fj*CBMXXX_GH87_ contains two tryptophan residues, one from each of the first and last loops of the protein, in *Cp*CBMXXX_GH16_ the site contains a tyrosine and a tryptophan residue, both on the last loop, similar to the placement of two of the three tryptophans in the *Fj*CBMXXX_GH87_ β-site.

Since the two CBMXXX domains investigated in our study both contained only the β- and γ-site, we screened the members of the family for any apparent binding sites at the α-position using multiple sequence alignments of the same 250 proteins from the tree displayed in **Fig 5**. A smaller alignment of selected domains is shown in **Fig. 6**, which illustrates that there are indeed proteins with aromatic amino acids at the corresponding α-site, including proteins containing all three putative binding sites. The aromatic amino acid residues present at these sites are typically tryptophan, but all other aromatic amino acids, phenylalanine, tyrosine and histidine, also occur at the corresponding positions. Whether all sites in the unstudied domains are functional, and whether they facilitate the same binding affinities, remains to be elucidated. However, the large diversity of binding sites could potentially indicate diverse binding specificities, as is to be expected in domains that append such a varying distribution of catalytic domains.

**Figure 6.**
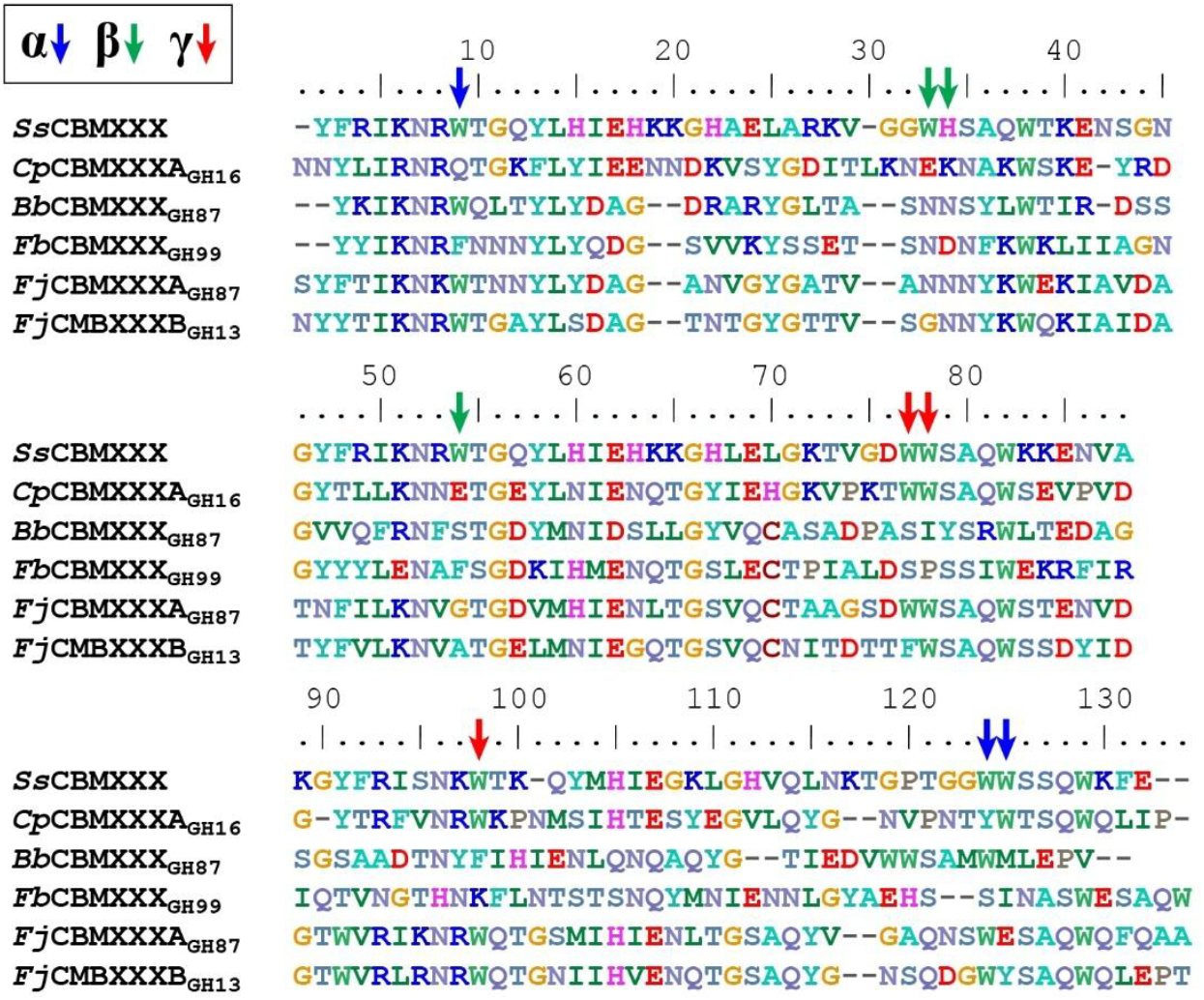
Multiple sequence alignment of selected members from CBMXXX. The proteins come from different bacteria, and their associated catalytic domains are indicated, where applicable: *Ss*CBMXXX from a *Sulfuvorum* sp. bacterium (CAA6815020.1), *Cp*CBMXXX_GH16_ from *Clostridium perfringens* (35), *Bb*CBMXXX_GH87_ from a *Bacteroidales* bacterium (MFN8205908.1,(36)), *Fv*CBMXXX_GH99_ from a *Flavobacteriaceae* bacterium (MDA7567898.1, (37)), and the two CBMXXX proteins from *Flavobacterium johnsoniae*, studied here. The positions of the binding residues are shaded and the respective binding site marked by arrows. Note that not all of the putative α-site binding residues align to the majority of the other sequences for *Bb*CBMXXX_GH87_ and *Fv*CBMXXX_GH99_.

### Comparison of CBMXXX to other β-trefoil CBM families

Based on sequence, CBMXXX is not homologous to other β-trefoil CBM families in CAZy. The sequence identity between CBMXXX domains and domains from either of CBM13, CBM42, and CBM92 is generally below 20 %, selected domains are compared in **Table S2**. Structurally, the β-strands in β-trefoils are conserved and provide the foundation for structural stability, while the loop regions are highly divergent among different families and provide variable functional roles (5). CBMXXX most closely resembles CBM13 and CBM92 in terms of structure, while CBM42 proteins have a more trianglular shape due to extended α-helices. However, the location of the binding residues differs in all families (**Fig. 7**). In CBMXXX, the binding residues originate from the loop close to the alpha helix and the loop following the two subsequent β-strands. In the case of the γ site, this means the binding residues originate from the beginning and end of the polypeptide sequence (marked by dark dots in **Fig. 7**). In both CBM13 and CBM42, the binding residues also originate from different loops, though in both cases these are only separated by one β-strand. In comparison, the binding residues in CBM92 are localized together, both structurally and sequence-wise, which could indicate a lower degree of binding variability in that family compared to CBMXXX, though evidence to substantiate this hypothesis is needed.

**Figure 7.**
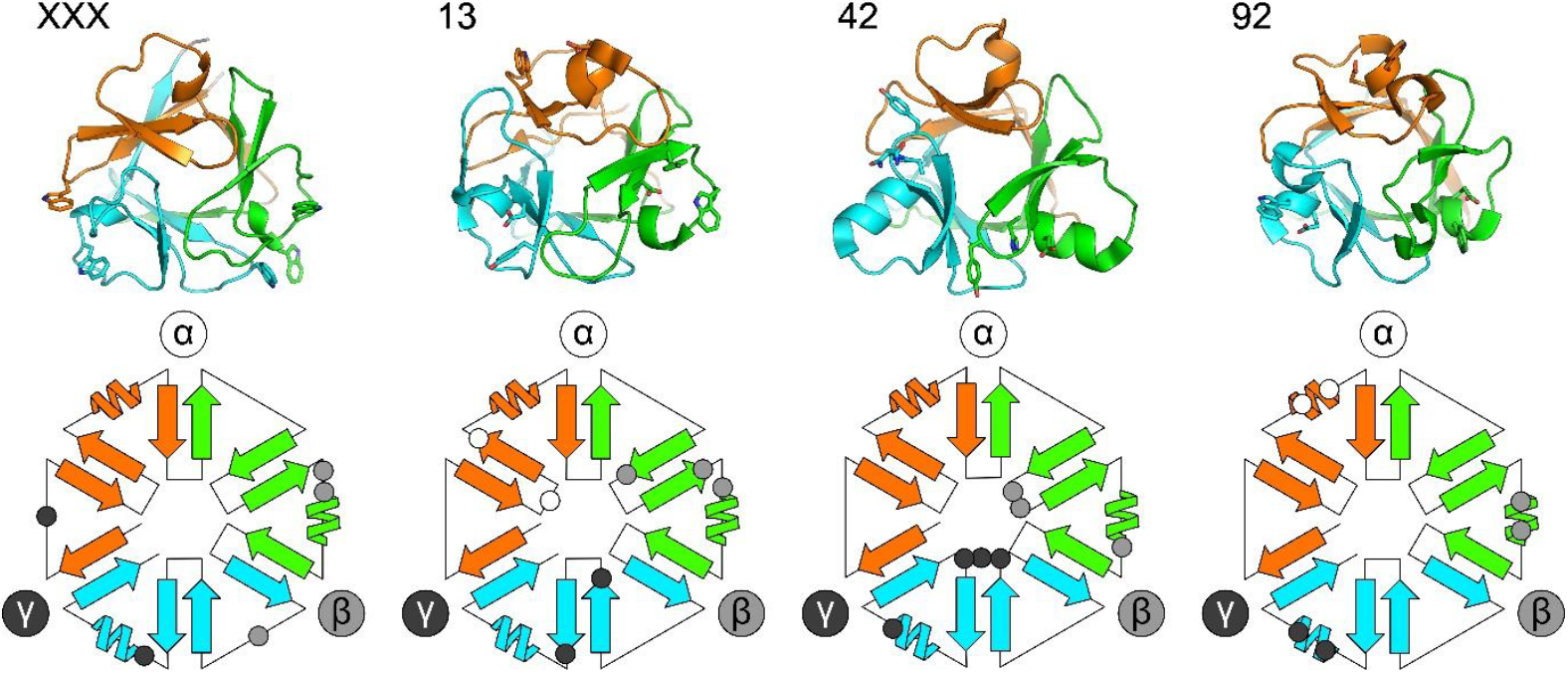
Structures and schematic representation of the CBM families with β-trefoil fold. Top row: from left to right, Alphfold model of *Fj*CBMXXX_GH87,_ crystal structures of representatives from CBM13 (PDB: 1KNM), CBM42 (PDB: 9NXI) and *Cp*CBM92 (PDB: 7ZON). Binding residues are shown as sticks. Bottom row: schematic representation of the same proteins. The binding residues’ positions are marked and color-coded based on which of the binding sites they are found in. CBMXXX is the only family where the binding residues’ positions are “unordered” sequence-wise, as it is for the γ-site.

## Conclusion

In this study we investigate the function of a novel CBM domain, *Fj*CBMXXX_GH87,_ which binds insoluble α-1,3-glucan and alternan polysaccharides and enhances the activity of the full length *Fj*GH87 enzyme compared to the sole catalytic domain. Furthermore, the protein binds to nigero-, malto-, and isomaltooligosaccharides in solution, but does so with seemingly different linkage preferences compared to the insoluble glucans. This indicates that not only the identified binding sites are important for binding, but most likely also the surrounding structure, accomodating different polysaccharides with varying ability.

*Fj*CBMXXX_GH87_ is the founding member of a new CBM family, clearly different from the other families in CAZy with β-trefoil folds, CBM13, CBM42 and CBM92, both in terms of sequence similarity and binding features. CBMXXX is mainly appended to enzymes from GH87, GH13, GH16, and GH99. Based on the different associated enzymes, and a large diversity in binding site composition, CBMXXX can be expected to bind a range of different ligands, though deeper studies of various proteins would be required to verify this, including binding- and structural studies. Given its ability to recognize α-1,3-glucans, *Fj*CBMXXX_GH87_, or other proteins from CBMXXX, could be used as tools for enhancing enzyme efficiency or to specifically target α-1,3-glucans present in dental plaque or fungal cell walls for imaging or therapeutic purposes. We further propose that newly discovered and characterized members from this family should be named following the here used nomenclature, with a species identifier–CBMXXX–associated catalytic domain(s), as in *Fj*CBMXXX_GH87_, to facilitate description of both origin and expected biological roles.

## Acknowledgements

We would like to acknowledge Lake Shimer, at the University of Michigan Medical School, for support in ITC measurements.

