## Supplemental Tables S1-2, Supplemental Figures S1-S11 for "Characterization of an α-glucan-binding module from *Flavobacterium johnsoniae* as a founding member of carbohydrate-binding module family XXX"

### Supplementary tables

**Table S1.** Primers used for the amplification of genes used in this study and for the mutagenesis to generate protein variants. Bold text signifies overhangs for InFusion cloning into expression vectors, and lower-case letters indicate mutation sites.

| Primer name | Sequence |
|---|---|
| FjohX_F | **CTTCCAGGGCCATAGT**AGTACAGCTACAGGAAGTTATTTTACC |
| FjohX_R | **TGGTGGTGCTCGAGTCTA**TGCCGAAGCTGCCTGAAAC |
| TW56_FjGH13_full_F | **CTTCCAGGGCCATAGT**CAGGATCCTCCGCAATAC |
| TW57_FjGH13_full_X_R | **TGGTGGTGCTCGAGTCTA**AACAGTTGTTGGTTCTAATTG |
| TW58_FjGH13_cat_R | **TGGTGGTGCTCGAGTCTA**AGCCGGAGCATTTGTATTG |
| TW59_FjGH13_cbm26_F | **CTTCCAGGGCCATAGT**AATCCAAATACGAATTTTACAG |
| TW60_FjGH13_cbm26_R | **TGGTGGTGCTCGAGTCTA**AATACCCGTTCCCGGATTAG |
| TW61_FjGH13_X_F | **CTTCCAGGGCCATAGT**ACTTCTGGAGGAGGAAATTATTATAC |
| TW62_FjohX_W768A_F | CATTAAAAACAAAgcGACGAATAATTATTTATATG |
| TW63_FjohX_W768A_R | TTCGTCgcTTTGTTTTTAATGGTAAAATAAC |
| TW64_FjohX_Y774A_F | GAATAATTATTTAgcTGATGCAGGGGCTAATG |
| TW65_FjohX_Y774A_R | GCATCAgcTAAATAATTATTCGTCCATTTG |
| TW66_FjohX_Y791A_F | CAAATAACAATgcCAAATGGGAAAAAAATAGCTG |
| TW67_FjohX_Y791A_R | CATTTGgcATTGTTATTTGCAACAGTTG |
| TW68_FjohX_W832A_W833A_F | GATCTGACgcGgcGAGCGCACAATG |
| TW69_FjohX_W832A_W833A_R | GTGCGCTCgcCgcGTCAGATCCTGC |
| TW70_FjohX_W853A_F | CAAAAACAGAgcGCAGACAGGAAG |
| TW71_FjohX_W853A_R | GTCTGCgcTCTGTTTTTGATTCTTACC |
| TW72_FjohX_W877A_F | CAAAATAGCgcGGAAAGCGCACAATG |
| TW73_FjohX_W877A_R | CTTTCCgcGCTATTTTGAGCTCCTATATATTGAG |
| TW74_FjohX_W832A_F | GATCTGACgcGTGGAGCGCACAATG |
| TW75_FjohX_W832A_R | GCTCCACgcGTCAGATCCTGCTG |
| TW84_FjohX_W832A_F | GCAGGATCTGACgcGTGGAGCGCACAATGGTC |
| TW85_FjohX_W832A_R | GTGCGCTCCACgcGTCAGATCCTGCTGCTG |
| TW86_FjohX_W877A_F | GCTCAAAATAGCgcGGAAAGCGCACAATGG |
| TW87_FjohX_W877A_R | GTGCGCTTTCCgcGCTATTTTGAGCTCC |

**Table S2.** Percent identity matrix for selected CBMXXX, CBM13, CBM42 and CBM92 domains.

| | 7ZON_ CBM92 | 1KNM_ CBM13 | 9NXI_ CBM42 | *Ss*CBMXXX | *Cp*CBMXXX$_{GH16}$ | *Bb*CBMXXX$_{GH87}$ | *Fb*CBMXXX$_{GH99}$ | *Fj*CBMXXX$_{GH87}$ | *Fj*CMBXXX$_{GH13}$ |
|---|---|---|---|---|---|---|---|---|---|
| 7ZON_ CBM92 | 100.00 | 27.52 | 11.32 | 16.88 | 10.13 | 17.57 | 18.67 | 18.29 | 15.85 |
| 1KNM_ CBM13 | 27.52 | 100.00 | 9.09 | 10.67 | 9.33 | 16.90 | 16.67 | 15.19 | 17.72 |
| 9NXI_ CBM42 | 11.32 | 9.09 | 100.00 | 16.25 | 14.81 | 16.67 | 13.75 | 17.05 | 11.36 |
| *Ss*CBMXXX | 16.88 | 10.67 | 16.25 | 100.00 | 36.00 | 26.23 | 29.27 | 37.10 | 33.06 |
| *Cp*CBMXXX$_{GH16}$ | 10.13 | 9.33 | 14.81 | 36.00 | 100.00 | 26.02 | 28.46 | 36.80 | 42.40 |
| *Bb*CBMXXX$_{GH87}$ | 17.57 | 16.90 | 16.67 | 26.23 | 26.02 | 100.00 | 27.20 | 38.40 | 38.40 |
| *Fb*CBMXXX$_{GH99}$ | 18.67 | 16.67 | 13.75 | 29.27 | 28.46 | 27.20 | 100.00 | 37.01 | 30.71 |
| *Fj*CBMXXX$_{GH87}$ | 18.29 | 15.19 | 17.05 | 37.10 | 36.80 | 38.40 | 37.01 | 100.00 | 60.00 |
| *Fj*CMBXXX$_{GH13}$ | 15.85 | 17.72 | 11.36 | 33.06 | 42.40 | 38.40 | 30.71 | 60.00 | 100.00 |

### Supplementary figures

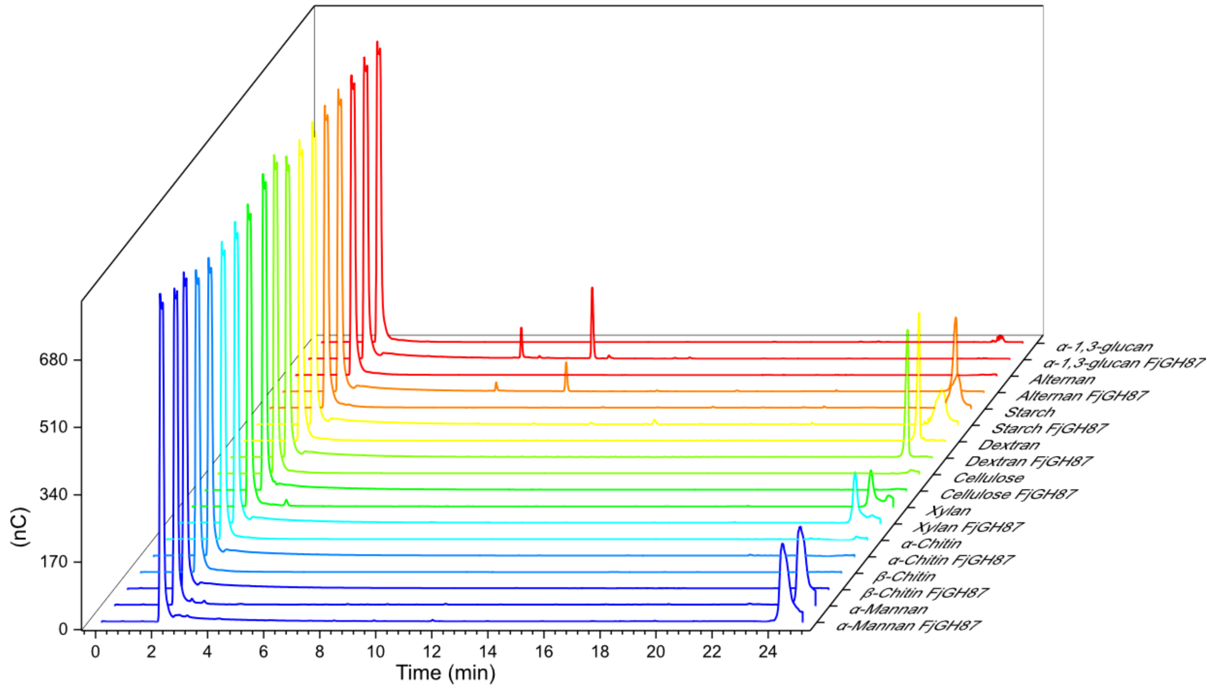


**Figure S1.** HPAEC-PAD analysis of the activity of *Fj*GH87 on polysaccharides together with no enzyme controls. Degradation of polysaccharides by *Fj*GH87 can only be seen for α-1,3-glucan and alternan. The main degradation products are nigerose and nigerotriose (see Fig. S2) for α-1,3-glucan and oligosaccharides of unknown size for alternan, but no glucose which would appear at a retention time around 3.5 min.

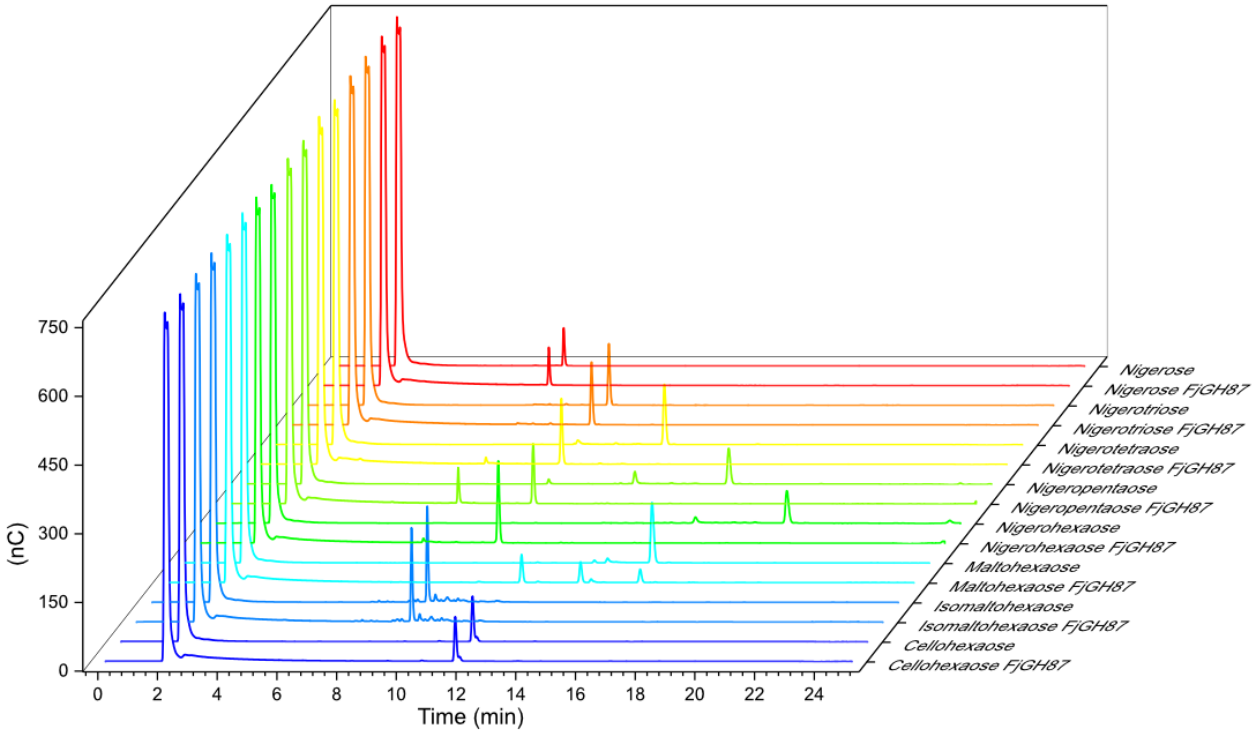


**Figure S2.** HPAEC-PAD analysis of the activity of *Fj*GH87 on oligosaccharides together with no enzyme controls. *Fj*GH87 cleaves nigerohexaose, nigeropentaose and nigerotetraose but not smaller oligosaccharides. It also has weak activity on maltohexaose but not isomaltohexose or cellohexaose.

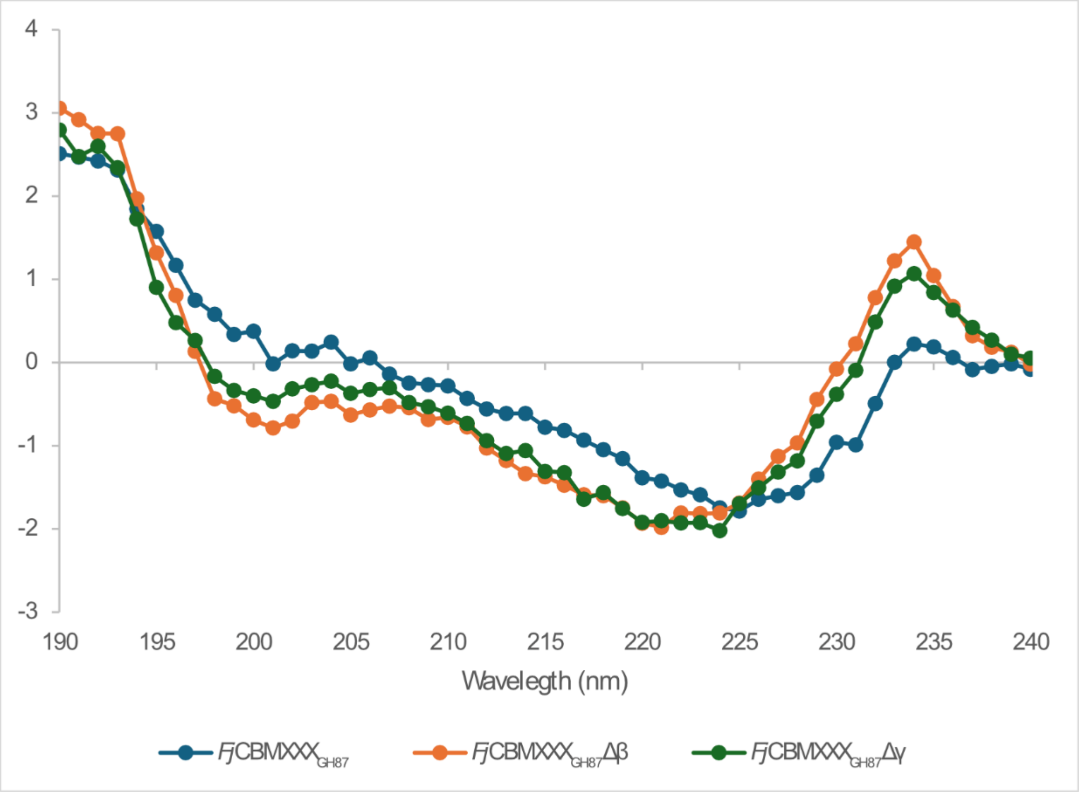


**Figure S3.** Circular dichroism of *Fj*CBMXXX$_{GH87}$, *Fj*CBMXXX$_{GH87}$-Δβ and *Fj*CBMXXX$_{87}$-Δγ.

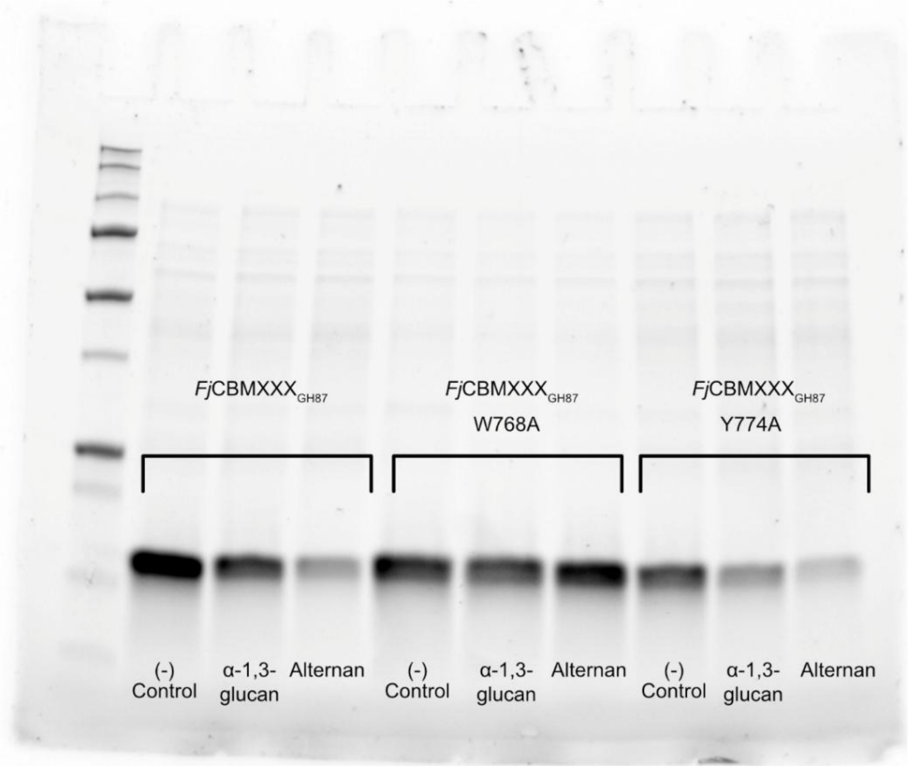


**Figure S4.** Pulldown assay of *Fj*CBMXXX$_{GH87}$ and variants. Left: *Fj*CBMXXX$_{GH87}$ (control, α-1,3-glucan, alternan), middle: *Fj*CBMXXX$_{GH87}$ W768A (control, α-1,3-glucan, alternan), right: *Fj*CBMXXX$_{GH87}$ Y774A (control, α-1,3-glucan, alternan).

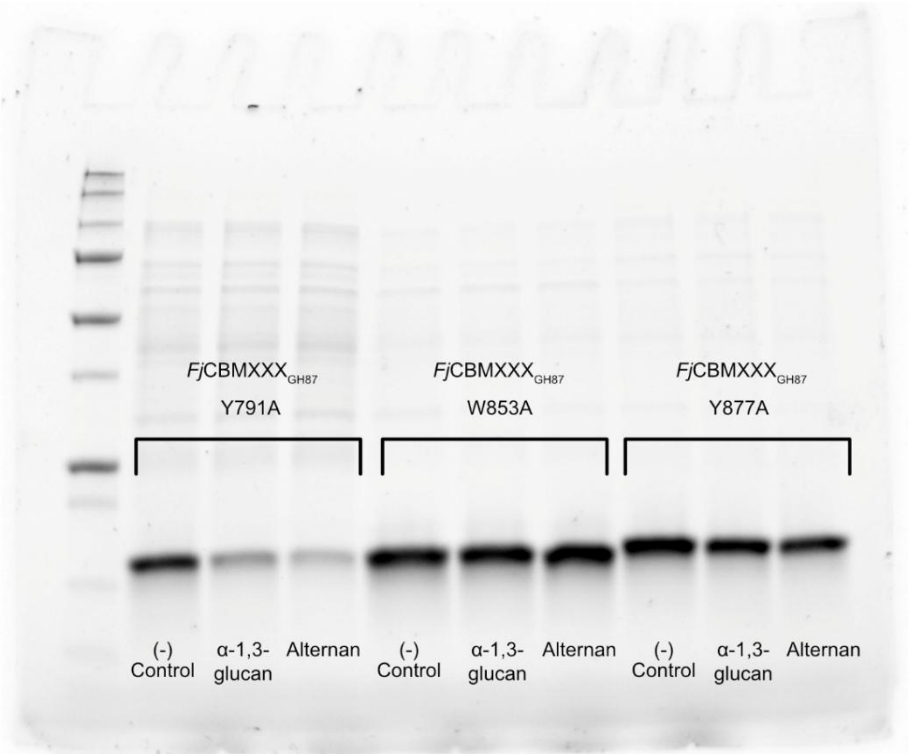


**Figure S5.** Pulldown assay of *Fj*CBMXXX$_{GH87}$ and variants. Left: *Fj*CBMXXX$_{GH87}$ Y791A (control, α-1,3-glucan, alternan), middle: *Fj*CBMXXX$_{GH87}$ W853A (control, α-1,3-glucan, alternan), right: *Fj*CBMXXX$_{GH87}$ W877A (control, α-1,3-glucan, alternan).

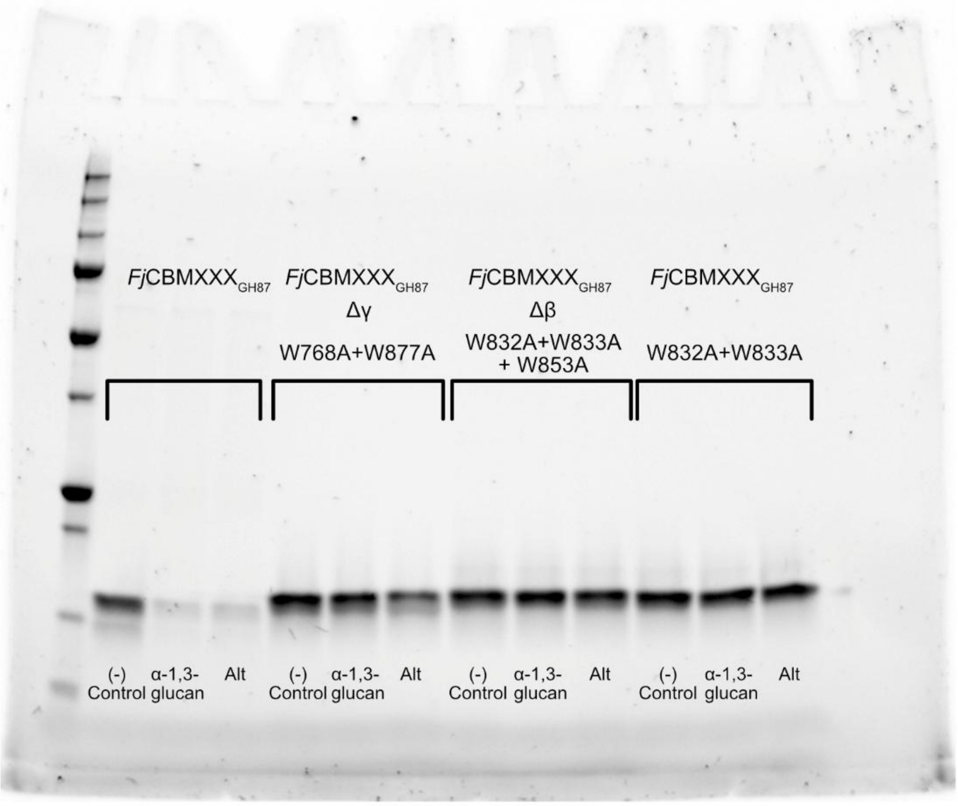


**Figure S6.** Pulldown assay of *Fj*CBMXXX$_{GH87}$ and variants. *Fj*CBMXXX$_{GH87}$ (control, α-1,3-glucan, alternan), *Fj*CBMXXX$_{GH87}$-Δγ (W768A + W877A; control, α-1,3-glucan, alternan),

*Fj*CBMXXX$_{GH87}$-Δβ (W832A+W833A+W853A; control, α-1,3-glucan, alternan), and *Fj*CBMXXX$_{GH87}$ partial β-site variant (W832A+W833A; control, α-1,3-glucan, alternan).

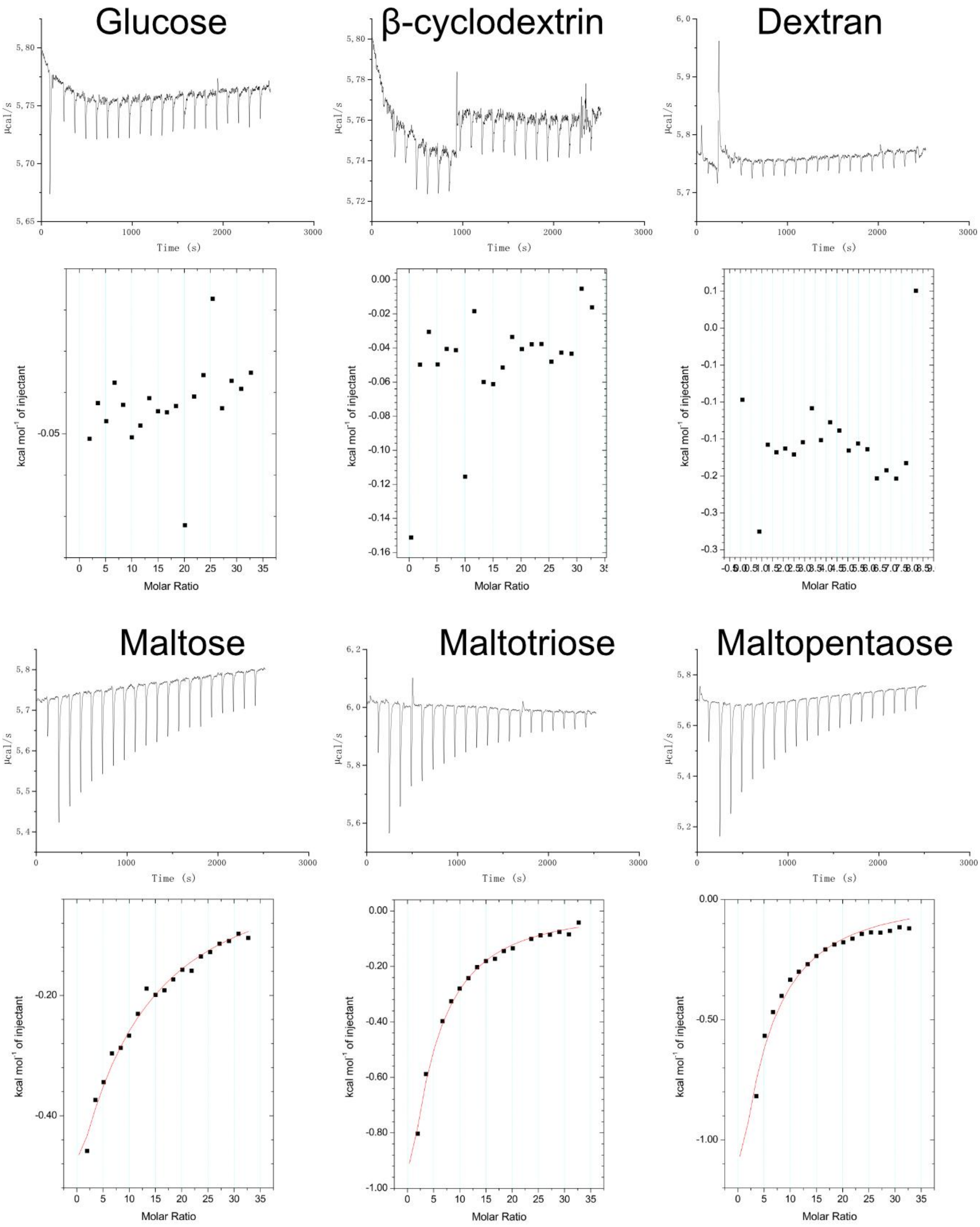

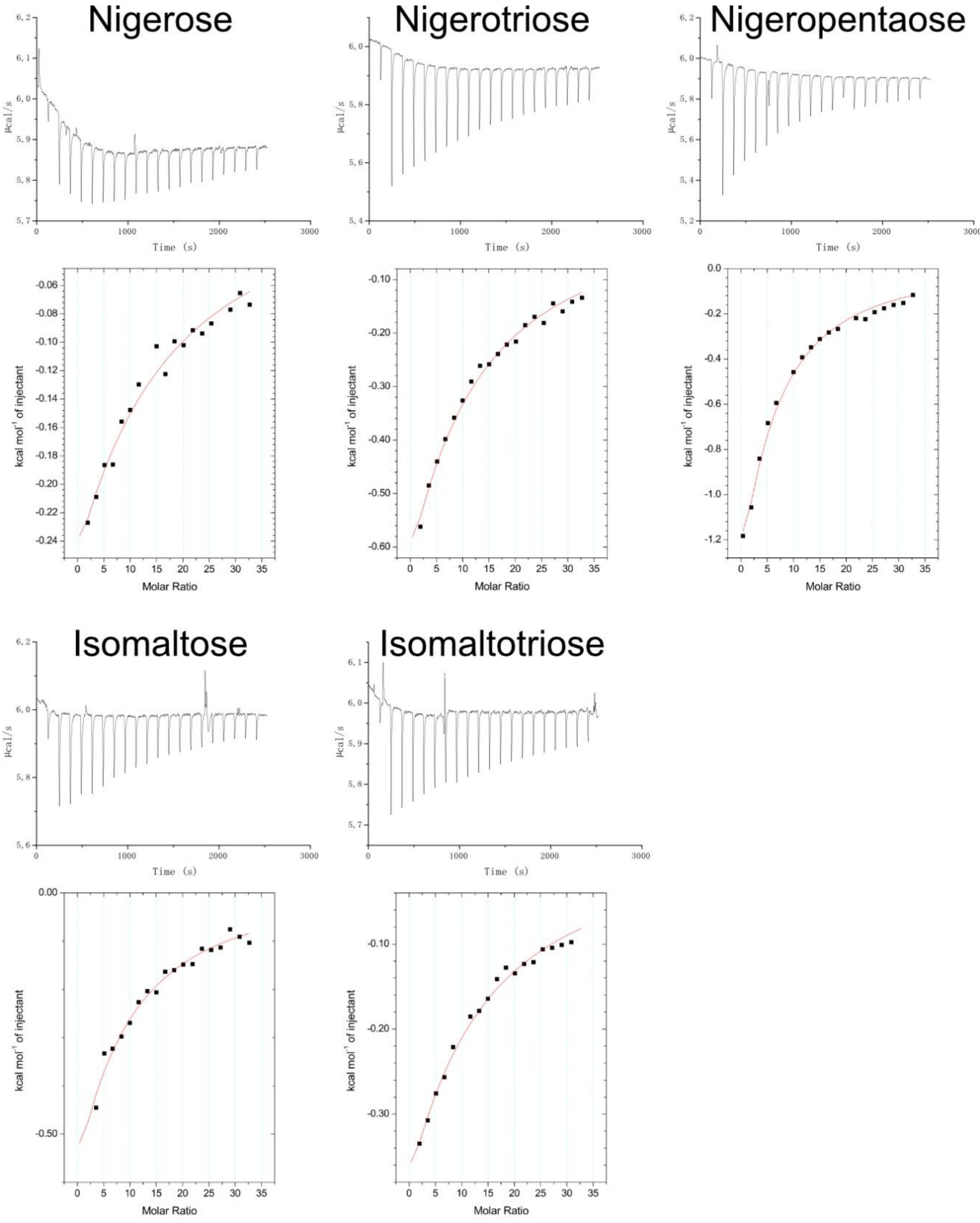


**Figure S7.** Representative graphs of the ITC measurements of *Fj*CBMXXX$_{GH87}$ with the different ligands glucose, β-cyclodextrin, dextran, maltose, maltotriose, maltopentaose, nigerose, nigerotriose, nigeropentaose, isomaltose, and isomaltotriose.

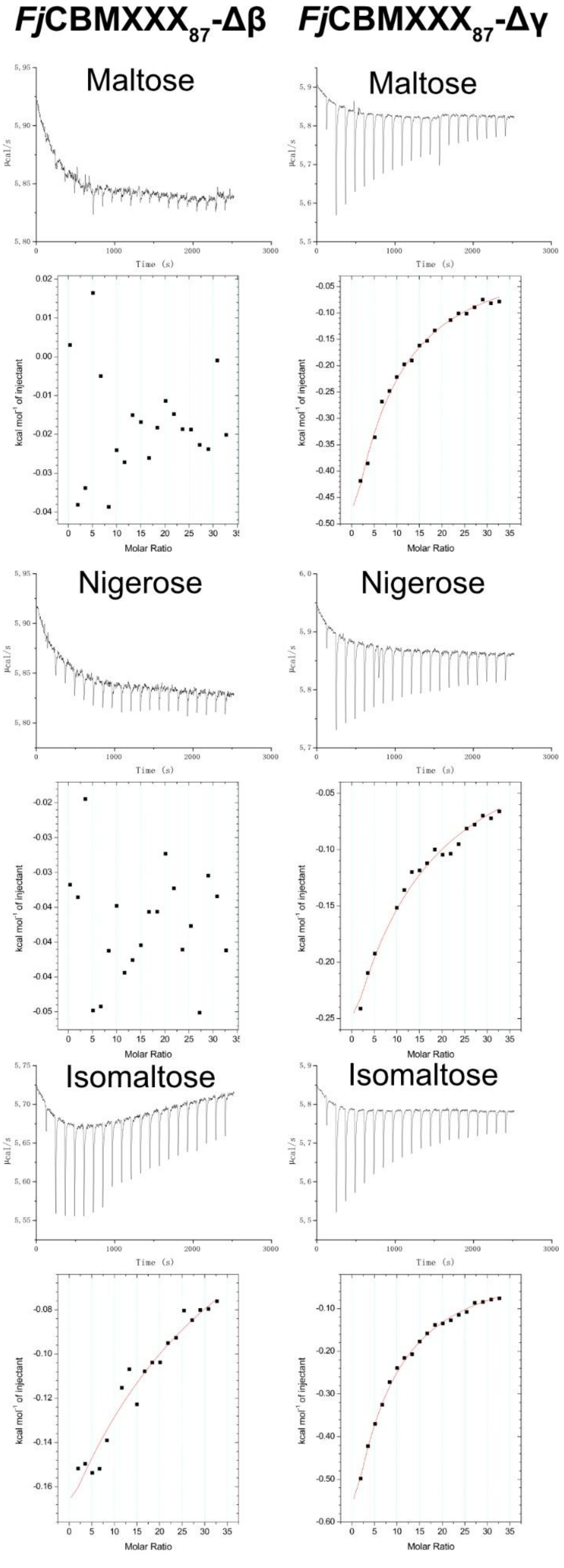


**Figure S8.** Representative graphs of the ITC measurements of the substitution variants *Fj*CBMXXX$_{87}$-Δβ and *Fj*CBMXXX$_{GH87}$-Δγ, with the ligands maltose, nigerose, and isomaltose.

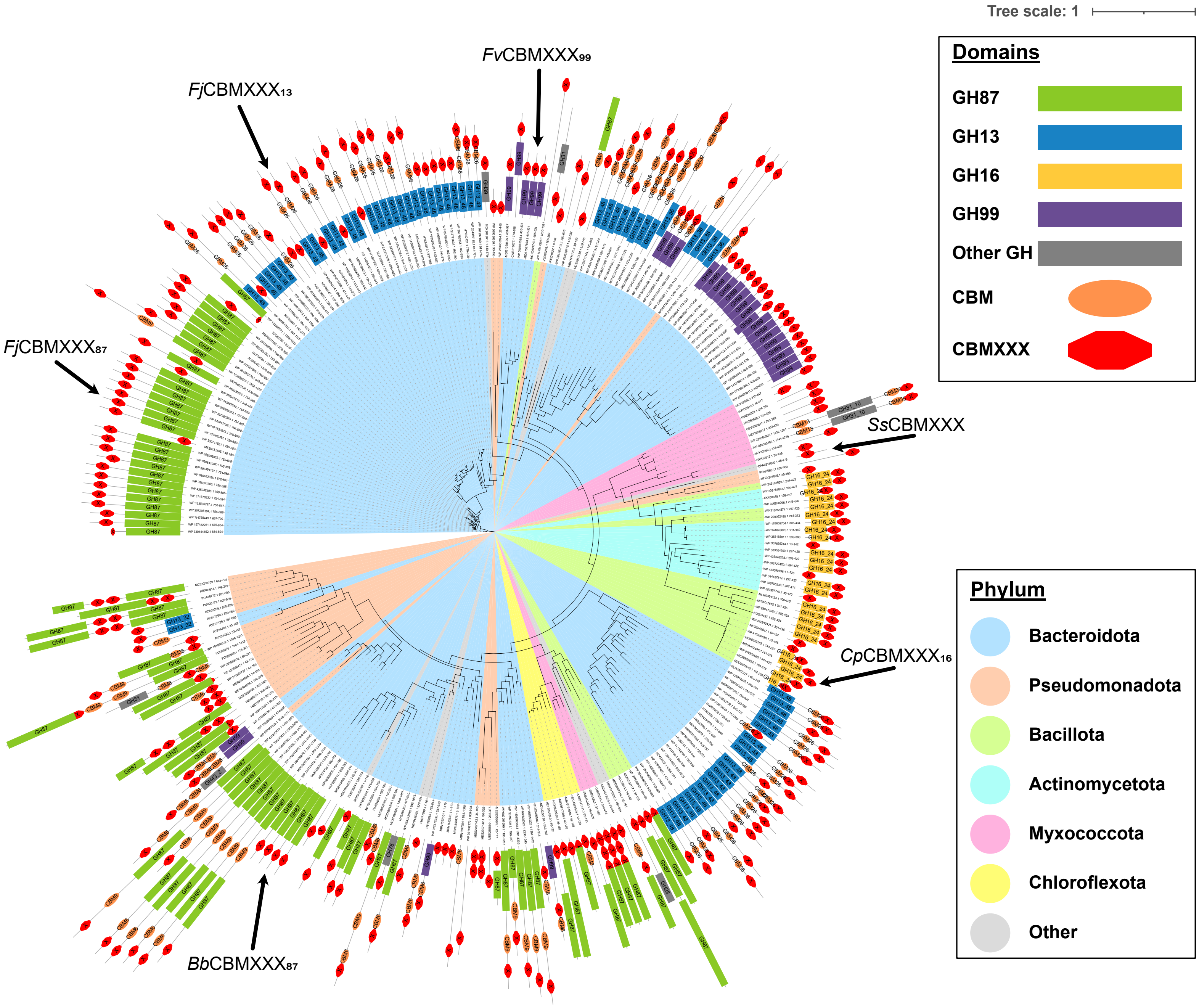


**Figure S9.** Phylogenetic tree of CBMXXX, with appended enzymes and other modules shown, as in Figure 5 in the main text. The GenBank IDs are shown for each entry, at the tip of the branches.

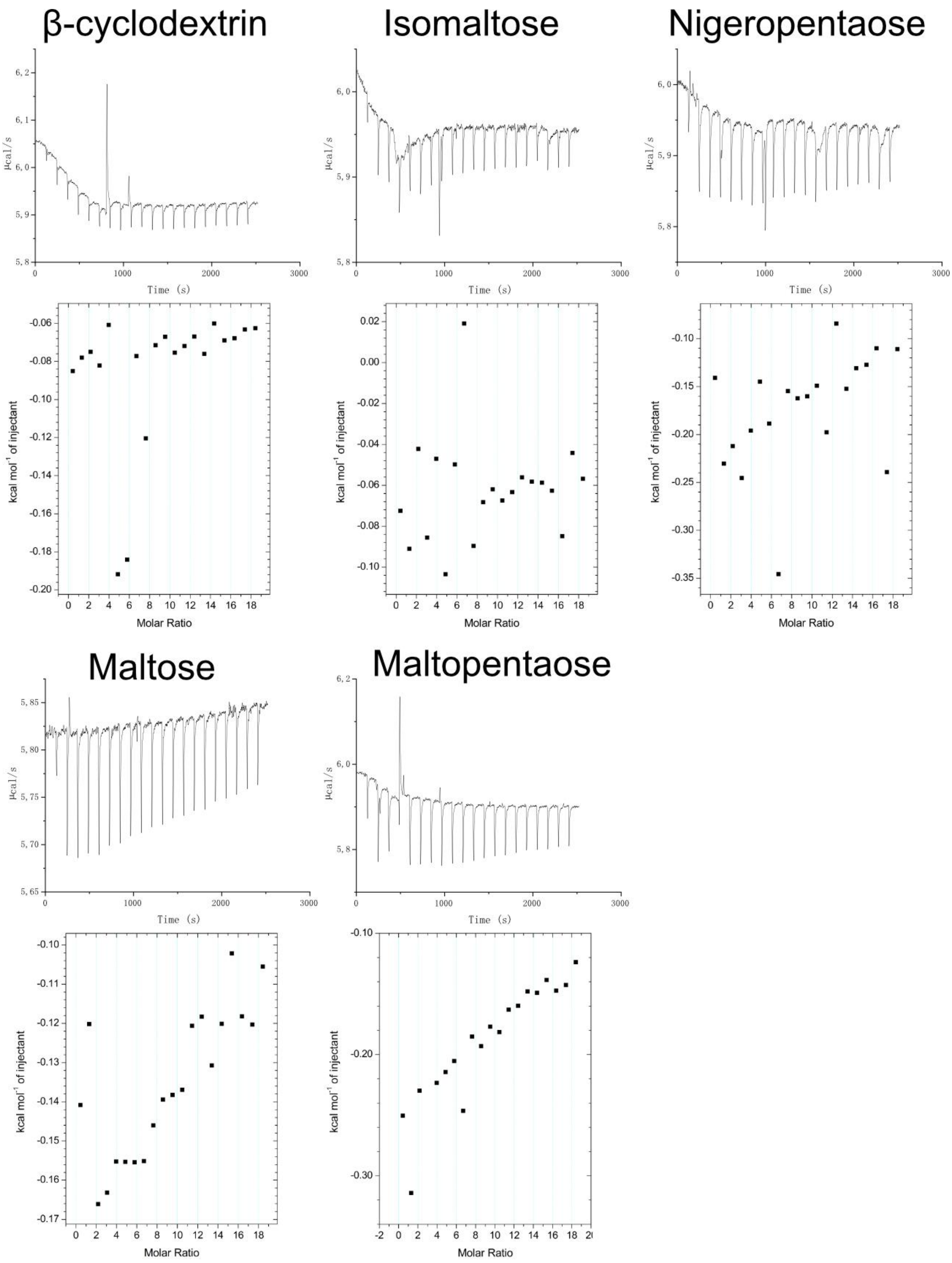


**Figure S10**. Graphs of the ITC measurements of *Fj*CBMXXX$_{GH13}$ with the ligands β-cyclodextrin, isomaltose, maltose, maltopentaose, and nigeropentaose.

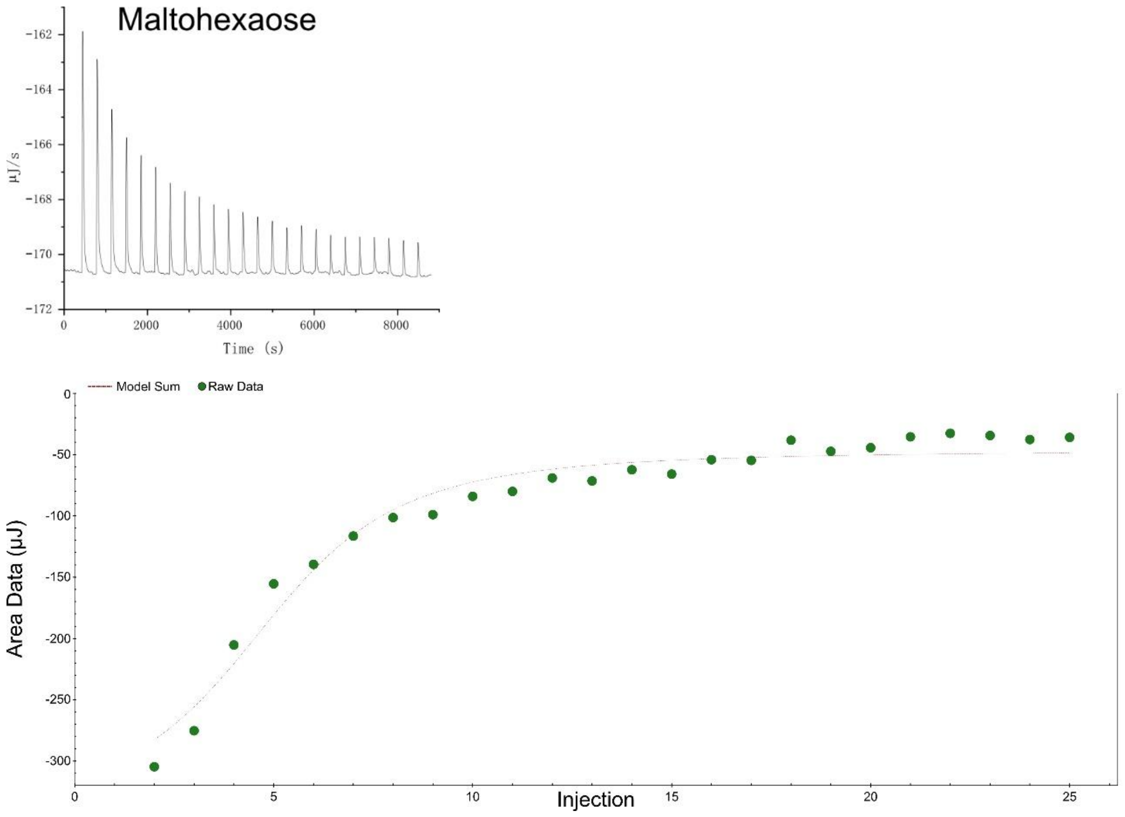


**Figure S11.** Graphs of the ITC measurements of *Fj*CBM26$_{GH13}$ with maltohexaose. Please observe that this experiment was done on a TA Instruments NanoITC, where y-axis unit is inverted.
